# Complexome profiling on the *lpa2* mutant reveals insights into PSII biogenesis and new PSII associated proteins

**DOI:** 10.1101/2021.01.04.425283

**Authors:** Benjamin Spaniol, Julia Lang, Benedikt Venn, Lara Schake, Frederik Sommer, Matthieu Mustas, Francis-André Wollman, Yves Choquet, Timo Mühlhaus, Michael Schroda

## Abstract

We have identified the homolog of LOW PSII ACCUMULATION 2 (LPA2) in *Chlamydomonas*. A *Chlamydomonas lpa2* mutant grew slower in low light and was hypersensitive to high light. PSII maximum quantum efficiency was reduced by 38%. Synthesis and stability of newly made PSII core subunits D1, D2, CP43, and CP47 were not impaired. Complexome profiling revealed that in the mutant CP43 was reduced to ∼23%, D1, D2, and CP47 to ∼30% of wild-type levels, while small PSII core subunits and components of the oxygen evolving complex were reduced at most by factor two. PSII supercomplexes, dimers, and monomers were reduced to 7%, 26%, and 60% of wild-type levels, while RC47 was increased ∼6-fold. Our data indicate that LPA2 acts at a step during PSII assembly without which PSII monomers and especially further assemblies become intrinsically unstable and prone to degradation. Levels of ATP synthase and LHCII were 29% and 27% higher in the mutant than in the wild type, whereas levels of the cytochrome *b*_*6*_*f* complex were unaltered. While the abundance of PSI core subunits and antennae hardly changed, LHCI antennae were more disconnected in the *lpa2* mutant, presumably as an adaptive response to reduce excitation of PSI. The disconnection of LHCA2,9 together with PSAH and PSAG was the prime response, but independent and additional disconnection of LHCA1,3-8 along with PSAK occurred as well. Finally, based on co-migration profiles, we identified three novel putative PSII associated proteins with potential roles in regulating PSII complex dynamics, assembly, and chlorophyll breakdown.

**One-sentence summary:** We provide evidence that the *Chlamydomonas* LPA2 homolog acts at a step in PSII biogenesis without which PSII monomers and further assemblies become unstable and prone to degradation.

## Introduction

Photosystem (PS) II is a light-driven water:plastoquinone oxidoreductase situated in the thylakoid membranes of cyanobacteria and chloroplasts. In land plants, the PSII core complex consists of the four large intrinsic subunits D1 (PsbA), D2 (PsbD), CP43 (PsbC), CP47 (PsbB), the 14 low-molecular-mass membrane spanning subunits PsbE, PsbF, PsbH, PsbI, PsbJ, PsbK, PsbL, PsbM, PsbTc, PsbW, PsbX, PsbY, PsbZ, and Psb30 and the five extrinsic subunits PsbO, PsbP, PsbQ, PsbR, and PsbTn (Pagliano et al., 2013; Wei et al., 2016; van Bezouwen et al., 2017). The latter stabilize and shield the Mn_4_CaO_5_ cluster and are attached to the lumenal surface of the core complex to form the oxygen evolving complex (OEC). PSII core monomers assemble into dimers to which, at both sides, light harvesting proteins (LHCII) bind to form PSII supercomplexes. In land plants, a PSII dimer binds two each of the monomeric minor LHCII proteins CP24 (LHCB6), CP26 (LHCB5), and CP29 (LHCB4) in addition to up to four major LHCII heterotrimers (Caffarri et al., 2009; Kouril et al., 2011). In *Chlamydomonas reinhardtii*, lacking CP24, a PSII dimer binds two each of the CP26 and CP29 monomers as well as up to six major LHCII heterotrimers (Tokutsu et al., 2012).

The individual steps leading to the assembly of PSII core complexes are well understood (Komenda et al., 2012; Shi et al., 2012; Nickelsen and Rengstl, 2013; Lu, 2016; Plochinger et al., 2016). PSII assembly starts with the synthesis of the α- and β-subunits (PsbE and PsbF) of cytochrome *b*_*559*_, which can accumulate in the membrane in the absence of the D1 and D2 proteins and interacts with newly synthesized D2 (Morais et al., 1998; Muller and Eichacker, 1999; Komenda et al., 2004). A newly synthesized D1 precursor first interacts with PsbI formed ahead, followed by their assembly with the D2-cytochrome *b*_*559*_ complex into the reaction center (RC) (Dobakova et al., 2007). After proteolytic processing of the D1 precursor at its C-terminus (Anbudurai et al., 1994), the RC is complemented with CP47 and the low molecular mass subunits PsbH, PsbL, PsbM, PsbR, PsbTc, PsbX, and PsbY to form the RC47 (or CP43-free PSII monomer) intermediate (Rokka et al., 2005; Boehm et al., 2012). The addition of CP43 along with the small subunits PsbK, PsbZ, and Psb30 leads to the formation of PSII monomers (Sugimoto and Takahashi, 2003; Rokka et al., 2005; Boehm et al., 2011). During photoactivation, the Mn_4_CaO_5_ cluster is attached to the luminal side of PSII monomers, followed by the PsbO, PsbP, and PSBQ proteins in chloroplasts (Bricker et al., 2012). After dimerization and attachment of LHCII the assembly is complete and the supercomplex is transferred from stroma-exposed membranes to grana stacks (Tokutsu et al., 2012; van Bezouwen et al., 2017).

PSII assembly is facilitated by auxiliary factors that transiently bind to discrete assembly intermediates and are not part of the final complex. Many, but not all, of these auxiliary factors are conserved between cyanobacteria and chloroplasts (Nixon et al., 2010; Nickelsen and Rengstl, 2013). Auxiliary factors involved in early steps of PSII de novo assembly include the membrane protein insertase Alb3 (Ossenbühl et al., 2004; Göhre et al., 2006), the TPR protein LPA1 (PratA in *Synechocystis*) that loads early PSII precomplexes with Mn^2+^ (Peng et al., 2006; Stengel et al., 2012), or HCF136 (YCF48 in *Syncechocystis*) that facilitates formation of the RC complex (Meurer et al., 1998; Komenda et al., 2008). One factor involved in a later step of PSII assembly is LPA2, which was proposed to mediate efficient synthesis or stability of newly made CP43 (Ma et al., 2007; RETRACTION, 2016). Note here that the article was retracted because of extensive figure manipulations, while the authors stated that they still believe in the validity of their results. Other auxiliary factors associated with PSII are involved in regulating the dynamics of PSII assembly states. Examples are MET1/TEF30 that interacts with PSII monomers and facilitates PSII supercomplex formation (Bhuiyan et al., 2015; Muranaka et al., 2016), or PSB33/LIL8 that controls the dynamics of LHCII (Fristedt et al., 2015; Fristedt et al., 2017; Kato et al., 2017). PSII can undergo repair after photodamage of the D1 protein. Here, many of the de novo assembly factors are involved also in the repair cycle, while others are specific for repair (Lu, 2016; Theis and Schroda, 2016).

The analysis of PSII assembly and the factors involved was greatly facilitated by the technique of blue-native polyacrylamide gel electrophoresis (BN-PAGE), which allows separating membrane protein complexes according to their size (Schagger and von Jagow, 1991; Schagger et al., 1994). A follow-up technique is complexome profiling, where a lane of a BN gel is cut along the gradient into dozens of even slices and proteins therein are identified and quantified by mass spectrometry after tryptic in-gel digestion (Heide et al., 2012; Heide and Wittig, 2013). By this, migration profiles are obtained for hundreds of proteins and proteins with similar migration profiles are likely present in common protein complexes. This technique has been applied to several photosynthetic organisms, including *Arabidopsis thaliana, Physcomitrella patens, Chlamydomonas reinhardtii*, and five cyanobacterial species resulting in the Protein Co-Migration Database for photosynthetic organisms (PCoM) (Takabayashi et al., 2017).

In this study we have identified the *Chlamydomonas* homolog of LPA2 and employed the complexome profiling technique to thylakoids isolated from wild type and a *lpa2* mutant to get new insights into the function of LPA2 in PSII biogenesis. The distinct migration profiles of PSII core subunits in wild type and *lpa2* mutant allowed the identification of novel proteins potentially associated with PSII based on their co-migration with the PSII core subunits.

## Results

### The *lpa2* mutant accumulates hardly any PSII supercomplexes, shows impaired growth and is sensitive to high light intensities

Ma et al. stated that there is no gene encoding LPA2 in cyanobacteria and *Chlamydomonas* (Ma et al., 2007; RETRACTION, 2016). To test this, we performed database searches and could confirm the absence of genes encoding LPA2 homologs in cyanobacteria but could find a gene encoding a putative LPA2 homolog in *Chlamydomonas*. We analyzed LPA2 homologs from land plants, moss and green algae and found that they all share properties such as a chloroplast transit peptide followed by a sequence with a twin-arginine motif, and a highly conserved region with two predicted transmembrane domains (Figure 1A).

**Figure 1.**
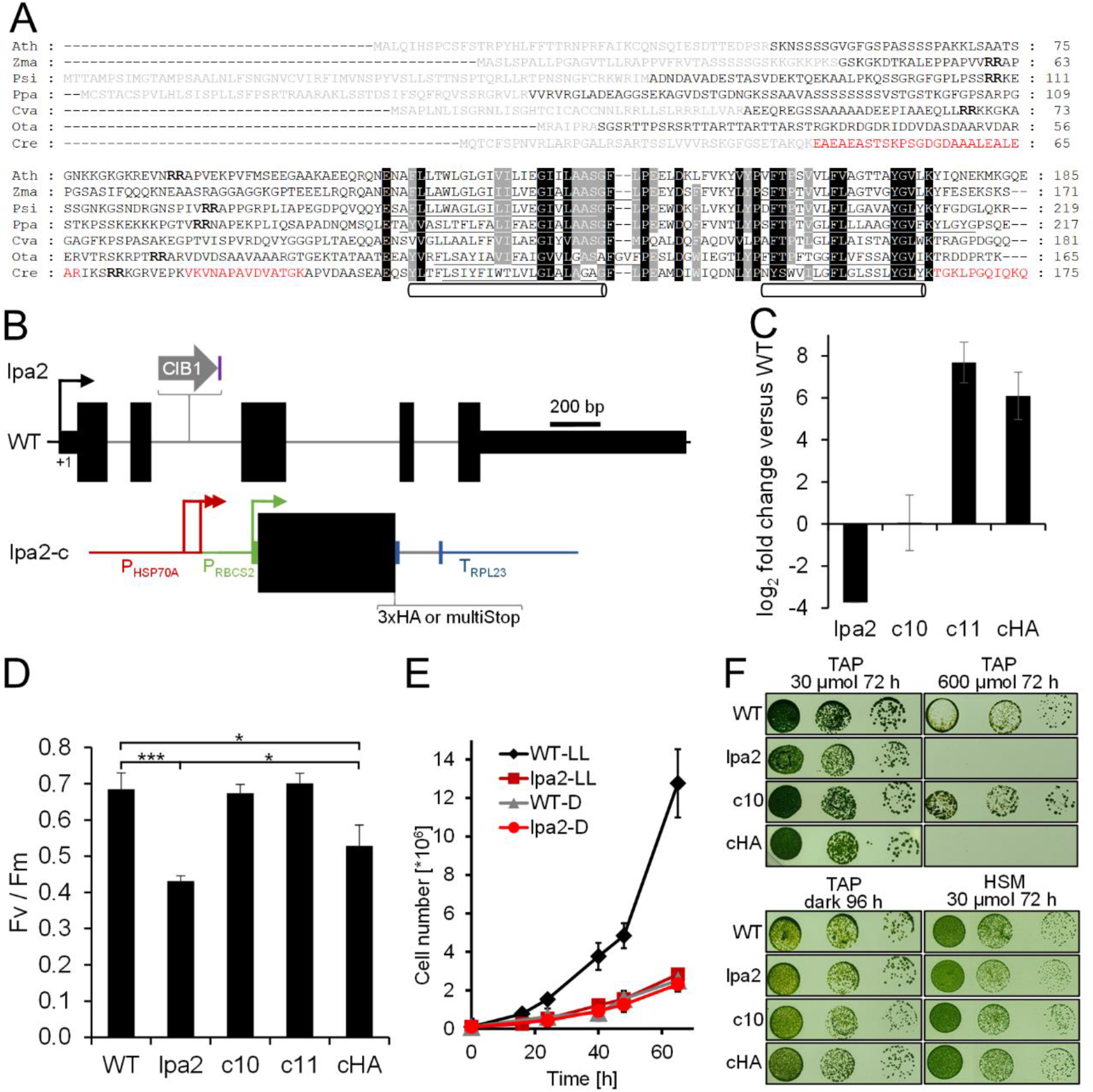
Growth and chlorophyll fluorescence phenotypes of the *lpa2* mutant compared to wild type and complemented lines. A, Alignment of LPA2 amino acid sequences from different organisms. Predicted chloroplast transit peptides are shown in grey, predicted transmembrane helices are underlined and indicated with pipes. Peptides identified by mass spectrometry are given in red letters, twin-arginines in bold letters. Ath – *Arabidopsis thaliana* (AT5G51545), Zma – *Zea mays* (NP_001145487), Psi – *Picea sitchensis* (ABK23742), Ppa – *Physcomitrella patens* (XP_024366975), Cva – *Chlorella variabilis* (XP_005849843), Ota – *Ostreococcus tauri* (XP_003084445), Cre – *Chlamydomonas reinhardtii* (Cre02.g105650). B, Structure of the *Chlamydomonas LPA2* gene, insertion site of the CIB1 cassette in the *lpa2* mutant, and construct for complementation. Protein coding regions are drawn as black boxes, untranslated regions as bars, and introns and promoter regions as thin lines. Arrows indicate transcriptional start sites. The purple box indicates a 165-bp fragment derived from the 19th intron of gene Cre13.g573450 in reverse orientation that has integrated together with the CIB1 cassette. C, qRT-PCR analysis of *LPA2* transcript accumulation. Values are means from two independent biological replicates and indicate *LPA2* transcript levels in the *lpa2* mutant and two complemented lines expressing the *LPA2* cDNA without (c10, c11) or with a 3xHA coding region (cHA) relative to the wild type. Error bars indicate SD. D, Comparison of PSII maximum quantum efficiency (Fv/Fm). Shown are averages from 3-6 independent experiments, error bars indicate SD. Significant differences were assessed via T-test, (*** p<0.001, * p<0.05). E, Growth curves of wild type (WT) and *lpa2* mutant in low light (LL, 30 µmol photons m^-2^ s^-1^) and in the dark (D). Values are means from 3 independent experiments; error bars represent SD. F, Analysis of the growth of 10^4^ – 10^2^ spotted cells kept under the conditions indicated.

To test, whether the *Chlamydomonas* protein is a *bona-fide* LPA2 homolog, we ordered a mutant from the *Chlamydomonas* library project (CLiP) (Li et al., 2016) that contains the mutagenesis cassette in the second intron of the putative *LPA2* gene (Figure 1B). The integration site of the cassette was verified by PCR on genomic DNA from the mutant (Supplemental Figure 1). Sequencing of the amplified fragments revealed that the cassette had integrated together with a 165-bp fragment from a distant gene, probably derived from genomic DNA released from lysed cells (Zhang et al., 2014). Although the cassette had inserted into an intron, expression of the putative *LPA2* gene was reduced by more than 13-fold in the mutant compared to the wild type judged by qRT-PCR analysis (Figure 1C). The mutant exhibited a strongly reduced PSII maximum quantum efficiency compared to the wild type (0.43 versus 0.69, Figure 1D) and slower mixotrophic growth in liquid culture at low light intensities, while heterotrophic growth in the dark was indistinguishable from the wild type (Figure 1E). On solid medium, growth of the mutant under mixotrophic and autotrophic conditions in low light was only mildly impaired, while heterotrophic growth in the dark was unimpaired when compared to the wild type (Figure 1F). In contrast, the mutant was unable to grow in high light under mixotrophic conditions. As judged by light microscopy, there were no obvious morphological differences between mutant and wild-type cells (Supplemental Figure 2). Western blot analysis revealed that the accumulation of PSII core subunits D1, CP43, and CP47 was strongly reduced in the mutant compared to the wild type, while LHCII and core subunits of PSI, the cytochrome *b*_*6*_*f* complex, and the ATP synthase appeared to accumulate normally (Figure 2A). ^14^C-acetate pulse-chase labeling revealed no differences in translation rates or stability of newly synthesized chloroplast-encoded subunits of the major thylakoid membrane complexes between mutant and wild type (Figure 2B). However, BN-PAGE analysis showed an abolished accumulation of PSII complexes, particularly of PSII supercomplexes, in the mutant compared to the wild type (Figure 2D).

**Figure 2.**
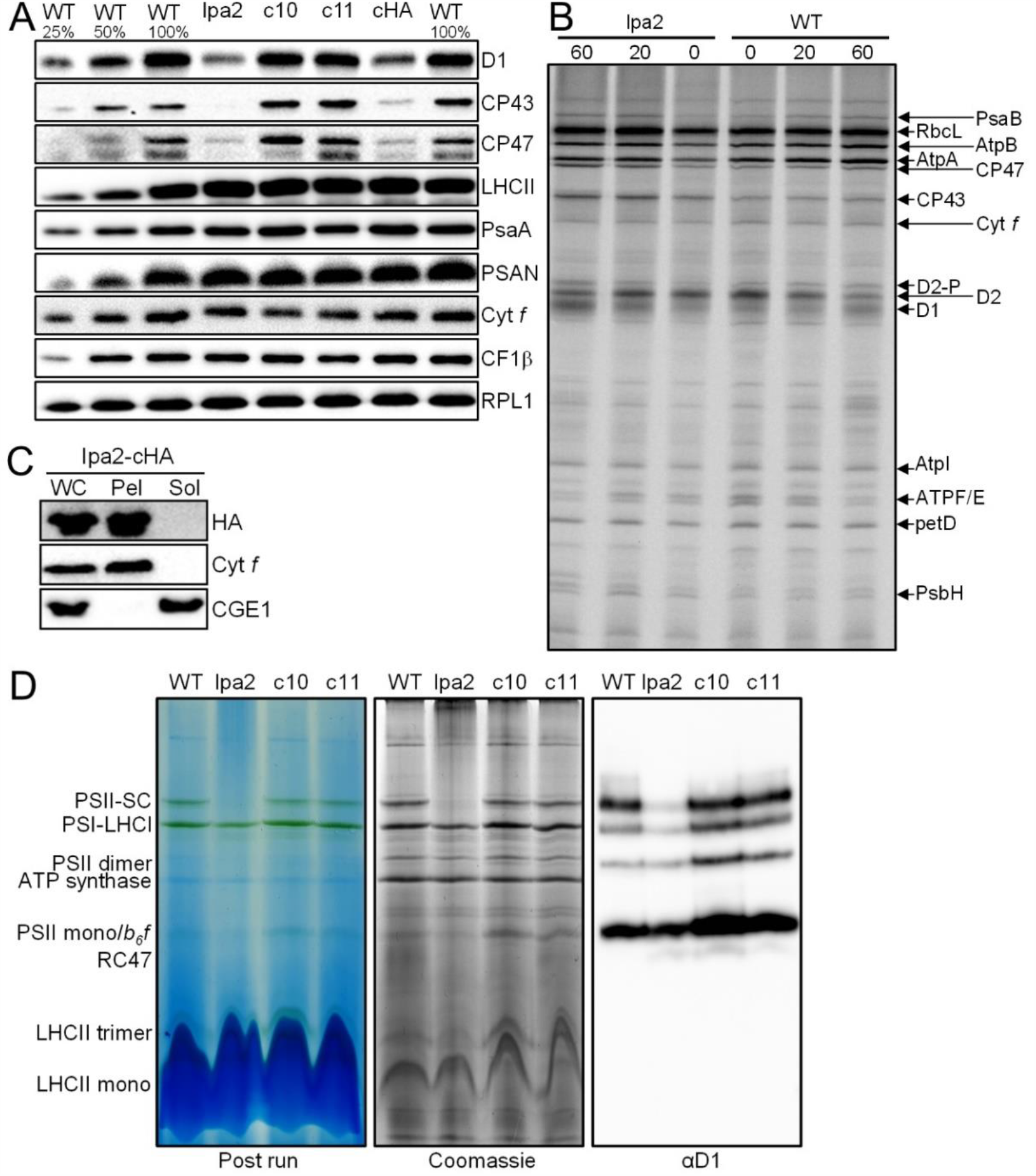
Analysis of protein accumulation, translation, localization, and complex assembly in *lpa2* mutant, wild type, and complemented lines. A, Immunoblot analysis of the accumulation of subunits of the major thylakoid membrane protein complexes. PSII – D1, CP43, CP47, LHCII; PSI – PsaA, PSAN; Cyt *b*_*6*_*f* complex – Cyt *f*; ATP synthase – CF1β. Ribosomal protein RPL1 serves as loading control. 10 µg of whole-cell proteins (100%) were analyzed. B, Comparison of translation and stability of newly translated proteins between wild type (WT) and *lpa2* mutant. Cells were labelled with ^14^C acetate for 7 min in the presence of cytosolic translation inhibitor cycloheximide (0) and chased with unlabelled acetate for 20 and 60 min. Proteins were separated on a 12-18 % SDS-urea gel and visualized by autoradiography. C, Crude fractionation of whole cells (WC) expressing LPA2-3xHA via freeze-thaw cycles and centrifugation into membrane enriched pellet (Pel) and soluble (Sol) fractions and analysis by immunoblotting with an anti-HA antibody. D, BN-PAGE analysis. Thylakoid membranes were prepared from wild type (WT), *lpa2* mutant, and complemented lines c10 and c11, solubilized with α-DDM, and proteins separated on a 4-15 % BN gel. Shown are pictures of the gel after the run, after staining with Coomassie, and after immunoblotting and detection with an antibody against D1. SC – supercomplexes.

To ensure that the observed phenotypes in the mutant were caused by the inactivation of the putative *LPA2* locus, we generated two constructs based on the *LPA2* cDNA and genetic parts from the Modular Cloning toolbox (Crozet et al., 2018) for complementation. The cDNA was driven by the *HSP70A-RBCS2* fusion promoter, terminated by the *RPL23* terminator, and translationally fused with parts encoding either multiple stop codons or a 3xHA epitope (Figure 1B). The two resulting level 1 modules were combined with an *aadA* cassette, transformed into the mutant, and resulting spectinomycin resistant transformants were screened for restored accumulation of the D1 protein (multi-stop construct) or accumulation of HA-tagged LPA2 protein (Supplemental Figure 3). Of 20 transformants screened for each construct, we identified four with fully restored D1 accumulation (multi stop) and one with detectable expression of the HA-tagged protein migrating in the SDS gel with an apparent molecular mass of 22.4 kDa. qRT-PCR revealed that multi-stop transformant c10 accumulated *LPA2* transcript to wild-type levels, while multi-stop transformant c11 accumulated it at ∼200-fold higher levels than the wild type (Figure 1C). The transformant harboring the construct encoding HA-tagged LPA2 accumulated *LPA2* transcripts at ∼70-fold higher levels than the wild type.

**Figure 3.**
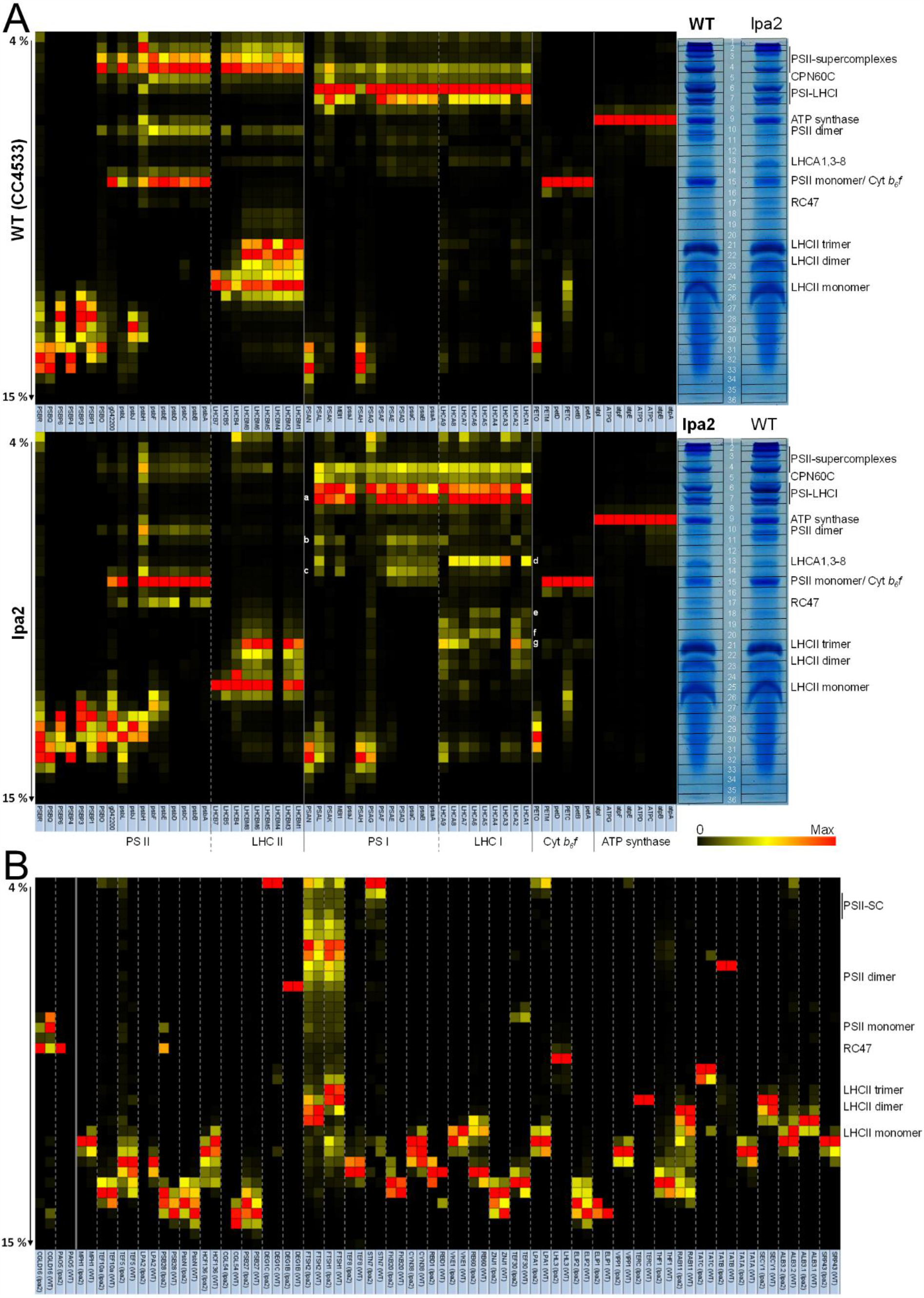
Complexome profiling on wild type and *lpa2* mutant. A, Heat map showing the BN-PAGE migration profiles of subunits of the major thylakoid membrane protein complexes of wild type (WT, top panel) and *lpa2* mutant (bottom panel). Values for each protein are derived from averaged peptide ion intensities from 3 biological replicates and are normalized to the gel slice with highest intensities. The BN-PAGE lane of one replicate from WT and *lpa2* mutant is shown with the excised band corresponding to the heat map row. White letters a-c and d-g indicate different assembly states of PSI core and LHCI antennae, respectively. The underlying data and the migration profiles for each protein are accessible in Supplemental Dataset 1. B, Heat map showing the BN-PAGE migration profiles of known (right from solid line) and putatively new (left from solid line) auxiliary factors involved in PSII biogenesis, repair, and the regulation of PSII complex dynamics in wild type and *lpa2* mutant. The accession numbers of all proteins are listed in Table S1.

While PSII maximum quantum efficiency was fully restored in multi-stop transformants c10 and c11, this was not the case for the transformant expressing HA-tagged LPA2 (cHA) (Figure 1D). Nevertheless, PSII maximum quantum efficiency in cHA was slightly but significantly higher than in the mutant. The slower growth and increased high light sensitivity phenotypes of the mutant were fully complemented in the multi-stop transformant c10 (and c11, not shown), while no complementation was observed for the cHA transformant (Figure 1F). Similarly, D1, CP43, and CP47 accumulated to wild-type levels in transformants c10 and c11, while in cHA these proteins accumulated only to slightly higher levels than in the mutant (Figure 2A). Apparently, the 3xHA epitope at the C-terminus interfered with full functionality of the LPA2 protein. Given the location of the two predicted transmembrane helices to the very C-terminus of LPA2, we wondered, whether the 3xHA tag disturbed proper membrane integration. To test this, we subjected cHA cells to freeze-thawing cycles and centrifugation for a crude separation of membrane and soluble proteins and detected transgenic LPA2 protein with an HA antibody. As shown in Figure 2C, the protein was detected exclusively in the pellet fraction. This indicates that the C-terminal 3xHA tag did not disturb proper membrane integration of LPA2, but apparently negatively impacted other aspects important for its functionality. The formation of PSII supercomplexes was fully restored in transformants c10 and c11 (Figure 2D).

In summary, inactivation of the putative *Chlamydomonas LPA2* gene resulted in reduced accumulation of PSII core subunits and PSII supercomplexes, reduced PSII maximum quantum efficiency, slower growth in low light in liquid cultures and increased sensitivity to high light. All these phenotypes could be complemented by expressing the wild-type protein, indicating that they are linked to the inactivated gene. As these phenotypes are very similar to those reported for the *Arabidopsis lpa2* mutant (Ma et al., 2007; RETRACTION, 2016), we can conclude that we have identified the *Chlamydomonas LPA2* gene.

### Comparing the complexome profiles of the *lpa2* mutant and wild type reveals novel insights into the role of LPA2 in PSII assembly and compensatory responses to cope with its absence

While most phenotypes observed for the *Arabidopsis* and *Chlamydomonas lpa2* mutants were very similar, this was not the case for the synthesis rate and stability of newly made CP43 (Ma et al., 2007; RETRACTION, 2016) (Figure 2B). Therefore, we reasoned that an in-depth comparison of the BN-PAGE migration profiles of thylakoid membrane proteins from wild type and *lpa2* mutant via complexome profiling (Heide et al., 2012) might reveal more insights into the function of LPA2. To this end we isolated thylakoid membranes from wild-type and *lpa2* mutant cells grown in low light (∼30 µmol photons m^-2^ s^-1^) in three biological replicates. Thylakoid membranes were solubilized with n-Dodecyl α-D-maltoside (α-DDM) and protein complexes separated on a 4-15 % BN gel (Supplemental Figure 4). Each gel lane was cut into 36 slices and the resulting 216 slices were subjected to tryptic in-gel digestion and LC-MS/MS analysis. In total, 1734 proteins were identified. Extracted ion chromatograms were used for protein quantification followed by normalization based on total ion intensities per lane (although equal amounts of protein were loaded, there are differences between the replicates that make normalization necessary (Supplemental Figure 4)). Ion intensity profiles for each protein along the BN gel run can be displayed from the Excel table in Supplemental Dataset 1. The migration profiles of all identified proteins of wild type and *lpa2* mutant, clustered according to their migration behavior, are shown in Supplemental Dataset 2 as heat maps. The profiles for proteins belonging to the major thylakoid membrane complexes from wild type and *lpa2* mutant are shown as heat maps in Figure 3. Missing subunits, such as PsbI, did not give rise to detectable peptides because peptides are too small, too large, too hydrophobic or contain posttranslational modifications other than methionine oxidation or N-acetylation.

**Figure 4.**
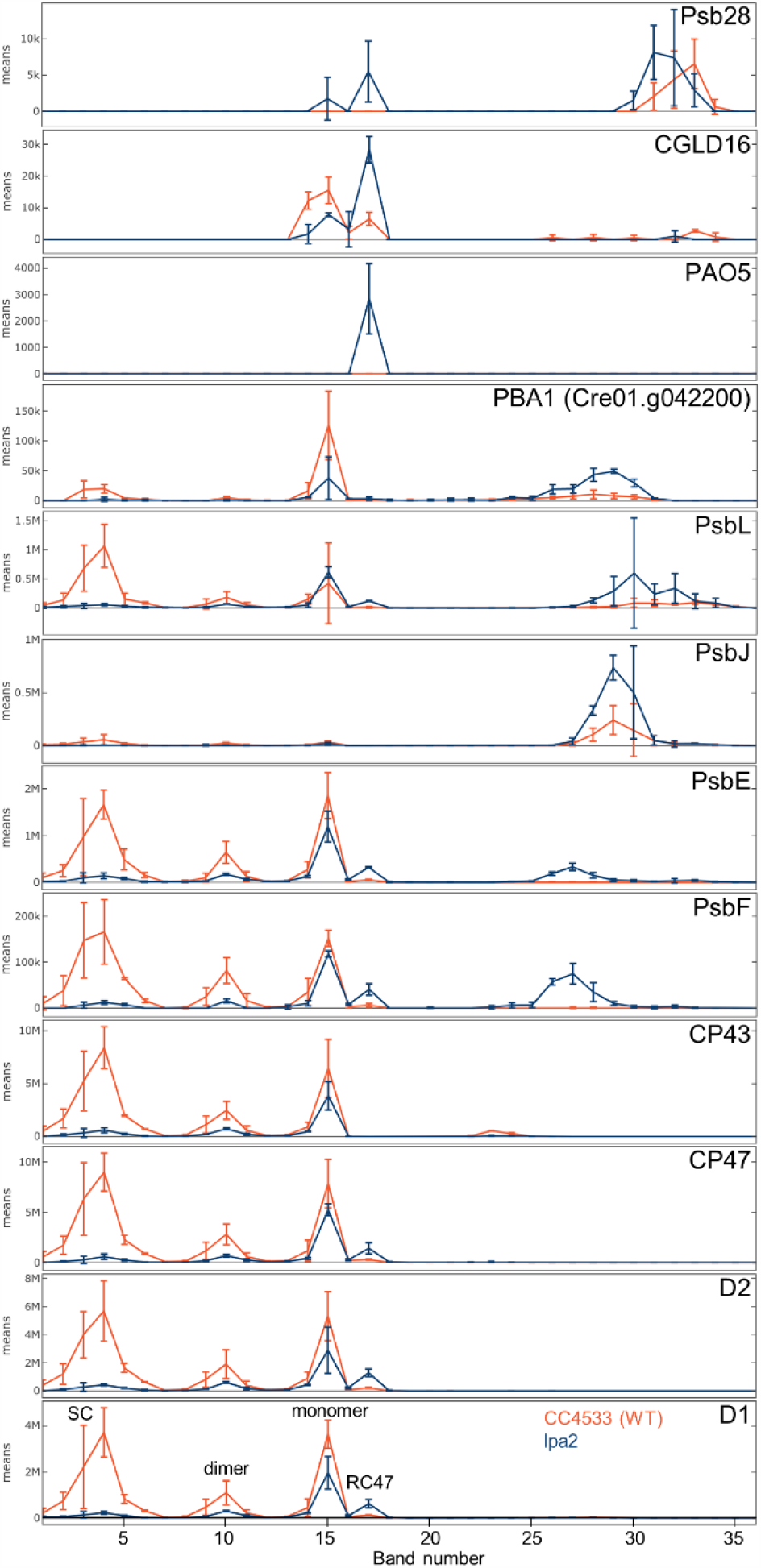
Comparison of BN-PAGE migration profiles of PSII core subunits and of putative novel proteins associated with PSII between wild type (red) and *lpa2* mutant (blue). Values for each protein are derived from averaged peptide ion intensities from 3 biological replicates. Error bars represent SD. Individual profiles from each replicate before and after normalization and statistical analyses can be accessed in Supplemental Dataset 1. SC – supercomplexes.

Eight subunits of the ATP synthase were identified and albeit their median abundance was∼29% higher in the *lpa2* mutant than in the wild type (Table 1), there was no difference in their migration patterns (Figure 3A). Six subunits of the cytochrome *b*_*6*_*f* complex were identified. They were about equally abundant in wild type and *lpa2* mutant and displayed the same migration behavior (Table 1, Figure 3A). As reported previously, PETO did not interact stably with other subunits of the complex (Takahashi et al., 2016). Interestingly, a substantial fraction of the Rieske iron-sulfur protein migrated as unassembled protein both in wild type and *lpa2* mutant, while all other subunits were quantitatively assembled into the complex.

**Table 1.** Ratio of subunit abundance between *lpa2* mutant and wild type. Values are based on the summed ion intensities in all gel bands of three biological replicates each of wild type and mutant.

| ATP synthase |  | Cytochrome *b<sub>6</sub>f* |  | Photosystem II |  | Photosystem I |  |
| --- | --- | --- | --- | --- | --- | --- | --- |
| Subunit | Ratio *lpa2*/WT | Subunit | Ratio *lpa2*/WT | Subunit | Ratio *lpa2*/WT | Subunit | Ratio *lpa2*/WT |
| AtpA | 1.27 | PetA | 1.01 | PsbA (D1) | 0.29 | PsaA | 0.93 |
| AtpB | 1.30 | PetB | 0.93 | PsbB (CP47) | 0.29 | PsaB | 0.71 |
| ATPC | 1.28 | PETC | 1.09 | PsbC (CP43) | 0.23 | PsaC | 0.88 |
| ATPD | 1.08 | PetD | 0.68 | PsbD (D2) | 0.30 | PSAD | 1.06 |
| AtpE | 1.52 | PETM | 1.52 | PsbE | 0.49 | PSAE | 1.15 |
| AtpF | 1.52 | PETO | 0.96 | PsbF | 0.55 | PSAF | 1.03 |
| ATPG | 1.28 | **Median** | **0.99** | PsbH | 0.67 | PSAG | 0.70 |
| AtpI | 1.72 |  |  | PsbJ | 2.17 | PSAH | 1.04 |
| **Median** | **1.29** |  |  | PsbL | 0.84 | PsaJ | 0.92 |
|  |  |  |  | PBA1 | 0.94 | PSAK | 1.35 |
|  |  |  |  | **Median** | **0.52** | PSAL | 0.82 |
|  |  |  |  | PSBO | 0.55 | PSAN | 1.25 |
|  |  |  |  | PSBP1 | 0.61 | **Median** | **0.98** |
|  |  |  |  | PSBP2 | 0.46 | LHCA1 | 1.10 |
|  |  |  |  | PSBP3 | 1.08 | LHCA2 | 1.19 |
|  |  |  |  | PSBP4 | 0.81 | LHCA3 | 0.88 |
|  |  |  |  | PSBP6 | 0.61 | LHCA4 | 0.96 |
|  |  |  |  | PSBQ | 0.54 | LHCA5 | 1.04 |
|  |  |  |  | PSBR | 0.50 | LHCA6 | 1.16 |
|  |  |  |  | **Median** | **0.58** | LHCA7 | 1.03 |
|  |  |  |  | LHCB4 | 1.20 | LHCA8 | 1.10 |
|  |  |  |  | LHCB5 | 1.45 | LHCA9 | 1.25 |
|  |  |  |  | LHCB7 | 1.79 | **Median** | **1.10** |
|  |  |  |  | LHCBM1 | 1.41 |  |  |
|  |  |  |  | LHCBM3 | 1.25 |  |  |
|  |  |  |  | LHCBM5 | 1.27 |  |  |
|  |  |  |  | LHCBM6 | 1.27 |  |  |
|  |  |  |  | LHCBM8 | 1.20 |  |  |
|  |  |  |  | LHCBM9 | 0.94 |  |  |
|  |  |  |  | **Median** | **1.27** |  |  |

In contrast to ATP synthase and cytochrome *b*_*6*_*f* complex, the composition of PSI complexes showed marked differences between wild type and mutant. The median abundance of PSI core subunits was unchanged and that of LHCI antennae only slightly higher in the mutant than in the wild type (Table 1, Figure 3A). However, LHCI antennae were much more disconnected in the *lpa2* mutant, with LHCA1 and LHCA3-8 still forming a common complex that was even visible in the BN-PAGE gel, in addition to smaller assemblies (d-f in Figure 3A). LHCA2 and LHCA9 were not present in this complex and accumulated only as smaller assemblies (g in Figure 3A as the major one). As a result of the disconnected antennae, more PSI core complexes were observed at smaller apparent molecular mass in the mutant compared to the wild type (a-c in Figure 3A). Moreover, in the mutant, more of the small PSI subunits PSAG, PSAL, PSAK, and PSAJ in this order were disconnected from the core complex and accumulated as unassembled subunits. Most PSAH and all PSAN accumulated as unassembled subunits in both, *lpa2* mutant and wild type, presumably because they lost connection to the PSI core during sample preparation or electrophoresis.

As expected, the most dramatic change between *lpa2* mutant and wild type was at the level of PSII supercomplexes, which accumulated only to ∼7% of wild type levels in the mutant, as judged from the median abundance of the core subunits in the supercomplexes (Table 2; Figures 3A and 4). The abundance of dimers and monomers was reduced in the mutant as well, but not as marked as that of the supercomplexes (∼27% and ∼60% of wild type levels, respectively). In contrast, ∼6 times more RC47 accumulated in the mutant compared to the wild type, and large amounts of unassembled Cyt *b*_*559*_ (PsbE/F) accumulated only in the mutant. Also, much larger amounts of unassembled PsbJ and PsbL accumulated in the mutant compared to the wild type. We observed no RC, PsbI-D1, or PsbE/F-D2 assembly intermediates. Regarding the overall differences in abundance of PSII core subunits between mutant and wild type, the biggest difference was found for CP43, which accumulated only to 23% of wild-type levels, followed by CP47, D1, and D2 accumulating to ∼30% of wild-type levels in the *lpa2* mutant (Table 1). The small core subunits accumulated to between 49% (PsbE) and 84% (PsbL) of wild-type levels. In contrast, PsbJ was more than 2-fold more abundant in the *lpa2* mutant than in the wild type.

**Table 2.**
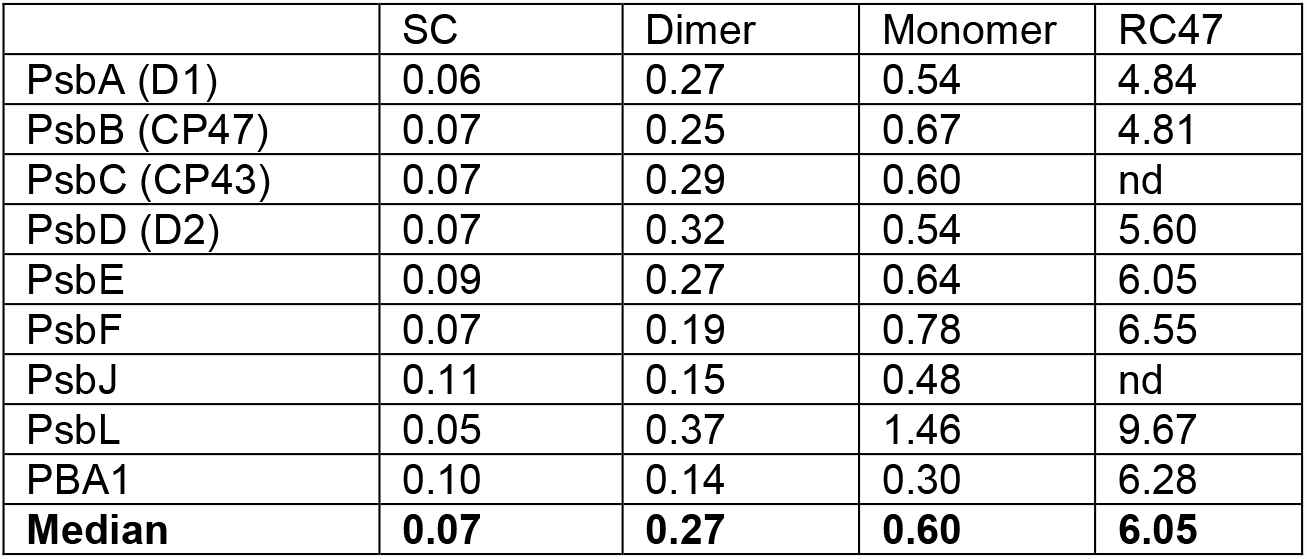
Ratio of subunit abundance in various PSII assembly states between *lpa2* mutant and wild type. Values are based on the summed ion intensities in bands 4 (supercomplexes, SC), 10 (dimers), 15 (monomers), and 17 (RC47) from three biological replicates each of wild type and mutant. nd – not detected.

Except for PSBO, all other subunits involved in stabilizing/shielding the Mn_4_CaO_5_ cluster were found to migrate as unassembled subunits in both, *lpa2* mutant and wild type, presumably because they got detached from PSII during sample preparation or electrophoresis. The median abundance of all subunits of the water-splitting complex reached 58% of wild-type levels. The median abundance of LHCII proteins was 27% higher in the mutant than in the wild type and therefore, since they could not be assembled into PSII supercomplexes, there was a large pool of unassembled LHCII trimers and monomers in the mutant. We could detect the recently identified LHCB7 protein with four transmembrane helices (Klimmek et al., 2006). However, in contrast to LHCB4 (CP29) and LHCB5 (CP26), LHCB7 accumulated only in the unassembled form (Figure 3A).

### The migration patterns of only few known PSII auxiliary factors differ between *lpa2* mutant and wild type

We reasoned that changes between the migration patterns of known auxiliary factors involved in PSII biogenesis, repair, and the regulation of PSII complex dynamics might reveal deeper insights into LPA2 function. Therefore, we started from a list of auxiliary factors compiled by Lu (2016) for *Arabidopsis* and searched for *Chlamydomonas* homologs that were detected in at least two replicates of mutant or wild type in the complexome profiling dataset (Supplemental Table 1). The migration profiles of the resulting 37 factors are displayed in the heat map shown in Figure 3B. Most of the factors were found to migrate in the low molecular mass region below LHCII trimers. Here, LPA2 and LPA19 (CGL54), a factor involved in the C-terminal processing of D1 during assembly (Wei et al., 2010), were completely missing in thylakoids from the mutant. Of the factors found in high molecular mass assemblies we found five to display differences between mutant and wild type based on data from at least two replicates: ALB3.2, LPA1, TEF5, TEF30, and PSB28. ALB3.2, which is required for the insertion of photosystem core proteins into the thylakoid membrane (Göhre et al., 2006), was found as unassembled protein and in a very large assembly hardly entering the gel. In the mutant, the balance between both forms was shifted to the unassembled protein. LPA1, an integral membrane protein required for proper PSII assembly (Peng et al., 2006), also migrated as unassembled protein and in a very large assembly. The latter was less abundant in the mutant. TEF5 (PSB33 or LIL8 in *Arabidopsis*), involved in regulating LHCII dynamics (Fristedt et al., 2015; Fristedt et al., 2017; Kato et al., 2017), was more abundant in the mutant and migrated in several higher molecular mass assemblies only in the mutant. TEF30 (MET1 in *Arabidopsis*) interacts with PSII monomers and facilitates PSII supercomplex formation (Bhuiyan et al., 2015; Muranaka et al., 2016). We found less TEF30 to migrate with PSII monomers in the *lpa2* mutant, in line with the lower amount of PSII monomers in the mutant. Psb28 has been found to associate with PSII monomers, RC47, and non-assembled CP47 in *Synechocystis* and was suggested to play roles during PSII repair and during PSII and PSI biogenesis, by mediating the incorporation of chlorophyll (Dobakova et al., 2009; Boehm et al., 2012; Sakata et al., 2013; Beckova et al., 2017). We found PSB28 to co-migrate with PSII monomers and RC47 only in the *lpa2* mutant.

### Identification of novel proteins potentially associated with PSI and PSII

We reasoned that the characteristic, distinct migration profiles of PSII and PSI subunits in wild type and *lpa2* mutant might enable the identification of novel proteins interacting with the photosystems. We therefore searched among the 1734 proteins identified in our complexome profiling dataset for proteins that meet the following four criteria: 1. Exhibit a migration profile resembling that of canonical PSI or PSII subunits in wild type and *lpa2* mutant. 2. Identified in at least two of the three replicates in wild type or mutant. 3. Contain a putative chloroplast transit peptide. 4. Are conserved in photosynthetic organisms. MBI1 met all four criteria to qualify as a protein potentially interacting with PSI. It co-migrated with PSI core subunits in wild type and *lpa2* mutant (Figure 3A). MBI1 is an OPR protein that functions as a maturation factor of the *psbI* transcript and appears to be conserved only in the Chlamydomonadales (Wang et al., 2015). Although MBI1 is a large protein (∼138 kDa), we identified only a single peptide (AEAEAQRRLGLGLR) with a comparably low identification score but good ion intensity (Supplemental Dataset 3). We therefore cannot rule out that the underlying fragmentation spectrum derives from a modified peptide of a PSI core subunit that is isobaric with the assigned MBI1 peptide and produces similar fragment ions. Hence, this finding should be taken with greatest care.

The gene product of Cre01.g042200 met all four criteria to qualify as a protein potentially interacting with PSII (Figure 3A, Figure 4). It co-migrated with PSII supercomplexes, dimers, RC47 and, most pronounced, with monomers. Its abundance in these complexes was reduced in the *lpa2* mutant compared to the wild type, while its unassembled form was more abundant in the mutant. The protein encoded by Cre01.g042200 contains 99 amino acids of which the N-terminal 36 were predicted to represent a chloroplast transit peptide (Figure 5A). The mature 63-amino acid protein has a molecular mass of 6.4 kDa and contains a predicted transmembrane helix. Two peptides were identified with high identification scores and good ion intensities (Supplemental Dataset 3). We found homologs only in members of the green algae, brown algae, diatoms, and Eustigmatophytes (Figure 5A). We named this protein PBA1 (putatively Photosystem B Associated 1).

**Figure 5.**
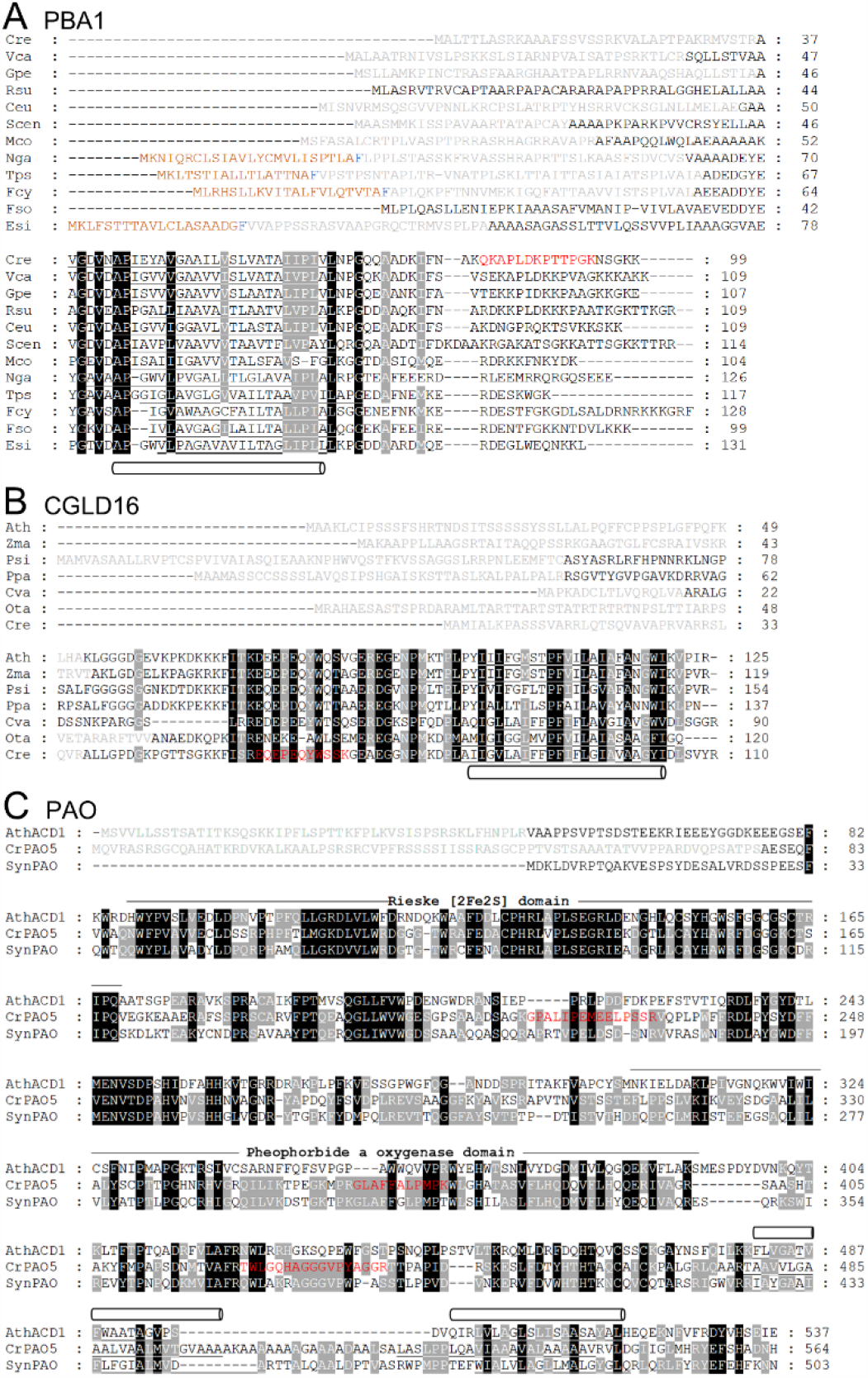
Alignment of amino acid sequences of putative novel PSII associated proteins from different organisms. Predicted chloroplast transit peptides are shown in grey and bipartite transit peptides from heterokonts are shown in brown with the conserved phenylalanine in blue (Kilian and Kroth, 2005). Predicted transmembrane helices are underlined and indicated with pipes. Peptides identified by mass spectrometry during complexome profiling are given in red letters. A, PBA1 homologs. Chlorophytes (green algae): Cre – *Chlamydomonas reinhardtii* (Cre01.g042200), Vca – *Volvox carteri* (Vocar.0001s0632), Gpe – *Gonium pectorale* (KXZ55153), Rsu – *Raphidocelis subcapitata* (GBF92771), Ceu – *Chlamydomonas eustigma* (GAX81760), Scen – *Scenedesmus sp*. (KAF6255734), Mco – *Micractinium conductrix* (PSC75504). Eustigmatophytes: Nga – *Nannochloropsis gaditana* (EWM21362). Bacillariophytes (diatoms): Tps – *Thalassiosira pseudonana* (XP_002292489), Fcy − *Fragilariopsis cylindrus* (OEU08446), Fso – *Fistulifera solaris* (GAX25330). Phaeophytes (brown algae): Esi – *Ectocarpus siliculosus* (CBN78173). B, CGLD16 homologs. Ath – *Arabidopsis thaliana* (AT2G05310), Zma – *Zea mays* (ACG27371), Psi – *Picea sitchensis* (ABR16811), Ppa – *Physcomitrella patens* (XP_024358733), Cva – *Chlorella variabilis* (XP_005848382), Ota – *Ostreococcus tauri* (XP_003078680), Cre – *Chlamydomonas reinhardtii* (Cre03.g151200). C, ACD1 from *Arabidopsis thaliana* (AtACD1, AT3G44880) compared to PAO5 homologs from *Chlamydomonas reinhardtii* (Cre06.g278245) and *Synechococcus sp*. PCC 7335 (SynPAO, EDX87484).

The accumulation of the PSII assembly intermediate RC47 to ∼6-fold higher levels in the *lpa2* mutant compared to the wild type (Table 2) opened the possibility to also identify novel auxiliary factors in PSII assembly that specifically interact with RC47. To this end, we searched in the complexome profiling dataset for proteins that accumulated in band 17 of the *lpa2* mutant at much higher levels than in the wild type and met criteria 2-4 introduced above. Three proteins met all criteria. The first is PSB28, which accumulated in band 15 (PSII monomers) and 17 (RC47) only in the *lpa2* mutant, and as free protein in mutant and wild type (Figures 3B and 4). The second protein, encoded by Cre03.g151200 (CGLD16), co-migrated with PSII monomers and RC47. In the *lpa2* mutant, less of this protein was present in the monomer band and more in the RC47 band compared to the wild type. CGLD16 is conserved in the green lineage and diatoms. The N-terminal 36 of the overall 110 amino-acids were predicted to form a chloroplast transit peptide, leaving a mature protein of 7.9 kDa that contains a predicted transmembrane domain at its C-terminus (Figure 5B). Only one peptide was identified for CGLD16 with high identification score and good ion intensity (Supplemental Dataset 3). The last protein, encoded by Cre06.g278245 (PAO5), co-migrated only with RC47 and only in the *lpa2* mutant. It contains a predicted chloroplast transit peptide, a Rieske [2Fe-S2] domain, and a pheophorbide a oxygenase (PAO) domain (Figure 5C). The mature protein has a molecular mass of 52 kDa. Three independent peptides were identified for PAO5 with rather low identification scores presumably due to the low precursor ion intensities (Supplemental Dataset 3). PAO5 is a member of a gene family with eight members in *Chlamydomonas*, while there are only three family members in *Arabidopsis thaliana* (Supplemental Figure 5). Interestingly, the closest homolog of PAO5 is *Chlamydomonas* PAO6 followed by cyanobacterial PAOs, while there is no close homolog of PAO5 in *Arabidopsis*.

In summary, complexome profiling allowed deep quantitative insights into defects in PSII assembly and possible adaptive responses of the *lpa2* mutant and allowed the identification of potentially novel proteins interacting with the photosystems.

## Discussion

### LPA2 from *Arabidopsis* and *Chlamydomonas* and mutants lacking LPA2 share many features

We report here on the identification of an LPA2 homolog in *Chlamydomonas* and its functional analysis. We found LPA2 to be conserved in the green lineage but to be absent in cyanobacteria (Figure 1A). All homologs share two predicted transmembrane helices at their C-termini, in accordance with the finding that both the *Arabidopsis* and *Chlamydomonas* protein behave like integral membrane proteins (Ma et al., 2007; RETRACTION, 2016) (Figure 2C). All LPA2 homologs contain a twin-arginine motif in the sequence following the predicted cleavage site of the chloroplast transit peptide (Figure 1A), but lack the highly hydrophobic residues at positions RR+2/3 and the A-X-A motif to qualify them as precursors for TAT pathway-mediated import into the thylakoid lumen (Cline, 2015). Accordingly, there is no processing of this sequence, as judged from the identification of two tryptic peptides in this sequence by mass spectrometry (Figure 1A; Supplemental Dataset 3). The twin-arginine motif might potentially aid in directing the protein to the thylakoid membrane for insertion.

The *Chlamydomonas lpa2* mutant shares many phenotypes with the *Arabidopsis lpa2* mutant (Ma et al., 2007; RETRACTION, 2016): both display slower growth (Figures 1E and 1F) and PSII maximum quantum efficiency is reduced to ∼65% of wild-type levels (Figure 1D). Both mutants accumulate slightly more ATP synthase (29% more in *Chlamydomonas*) (Table 1). Both accumulate PSII core subunits D1, D2, CP47, and CP43 to only ∼30% of wild-type levels, while subunits of the oxygen-evolving complex were more mildly reduced (Figure 2A; Table 1). For both, the most affected PSII assembly state is the supercomplexes, which are absent in the *Arabidopsis* mutant and accumulate to ∼7% of wild-type levels in the *Chlamydomonas* mutant (Figures 2D and 3A; Table 2). The *Arabidopsis lpa2* mutant contains 25% less PSI reaction center proteins, while their abundance is unaltered in the *Chlamydomonas* mutant (Table 1). However, reduced PSI acf tivity also in the *Chlamydomonas lpa2* mutant can be implied from the observation that the mutant had more disconnected LHCI antennae compared to the wild type (Figures 3A and 4), presumably to reduce excitation energy to PSI as a regulative adaptation to the low amounts of PSII. The disconnected LHCI antennae displayed characteristic migration profiles pointing to the disconnection of a major LHCI complex composed of LHCA1,3-8 (d in Figure 3A) and a minor one composed of LHCA2,9 (g in Figure 3A). The prominent LHCA1,3-8 complex appeared to disassemble into LHCA1,4-6, LHCA4-6, and LHCA1,7,8 subcomplexes (e-g in Figure 3A). This is in perfect agreement with a model proposed recently where LHCA1,3-8 bind at one side of the PSI core in two layers, with LHCA1,3,7,8 forming the inner and LHCA1,4-6 the outer layer; LHCA2,9 bind at the opposite side (Ozawa et al., 2018). A major PSI complex accumulating in the mutant, and to a lesser extent also in the wild type, specifically lacks LHCA2,9, PSAG and PSAH (a in Figure 3A). Again, this is in perfect agreement with the model that LHCA2,9 are flanked by PSAG and PSAH (Ozawa et al., 2018) and indicates that the uncoupling of these proteins is the prime response in the *lpa2* mutant to reduce the PSI antenna size. Nevertheless, a minor PSI complex contains LHCA2,9 but lacks LHCA1,3-8 (b in Figure 3A), indicating that the disconnection of LHCA2,9 must not necessarily precede the disconnection of LHCA1,3-8. Smaller PSI complexes presumably lacking LHCA1,3-8 also lack PSAK (c in Figure 3A), again in line with the proposed role of this subunit in flanking LHCA1,3-8 (Ozawa et al., 2018).

### Differences in phenotypes between *Arabidopsis* and *Chlamydomonas lpa2* mutants point to a role of LPA2 in a step required for the assembly of monomers into higher assembly states

A difference between the *Arabidopsis* and *Chlamydomonas* LPA2 proteins was that the protein was found to interact and to co-migrate with Alb3 in *Arabidopsis*, while we observed no co-migration between LPA2 and ALB3.1 or ALB3.2 in *Chlamydomonas* (Ma et al., 2007; RETRACTION, 2016) (Figure 3B). Moreover, the migration profiles in the lower molecular mass region of neither ALB3.1 nor ALB3.2 were altered in the *Chlamydomonas lpa2* mutant compared to the wild type. However, like in *Arabidopsis, Chlamydomonas* LPA2 was only found in assemblies of low molecular mass. There were also several phenotypical differences between the *Arabidopsis* and *Chlamydomonas lpa2* mutants. The *Chlamydomonas* mutant was highly sensitive to high light intensities, which was not reported for the *Arabidopsis* mutant (Figure 1F). While the *Arabidopsis* mutant accumulated slightly less LHCII than the wild type, we found LHCII levels in the *Chlamydomonas* mutant to be increased by ∼27% (Figure 2A; Table 1). Perhaps this larger pool of ‘free’ LHCII is the reason for the higher light sensitivity of the *Chlamydomonas* mutant? Moreover, the *Arabidopsis* mutant had slightly higher levels of the Cyt *b*_*6*_*f* complex than the wild type, while there was no change in the *Chlamydomonas* mutant.

The difference between the *Chlamydomonas* and *Arabidopsis lpa2* mutants that surprised us most was that the synthesis rate of CP43 and the stability of the newly made protein were not affected in the *Chlamydomonas* mutant, while CP43 synthesis rates were reduced to 25% of wild-type levels in the *Arabidopsis* mutant (with radioactive labeling times of 7 and 10 min, respectively) (Ma et al., 2007; RETRACTION, 2016) (Figure 2B). Since the *Chlamydomonas lpa2* mutant contained the mutagenesis cassette in the second intron of the *LPA2* gene (Figure 1B), it appears possible that this finding is due to residual LPA2 protein in the mutant. However, while LPA2 was detected with three peptides and good ion intensities in all replicates of the wild type, it was not detected in any of the replicates of the mutant (Supplemental Dataset 3). Hence, the unaffected synthesis rate and stability of newly made CP43 in the *Chlamydomonas lpa2* mutant argues against the previously proposed role of LPA2 in mediating efficient synthesis or stability of newly made CP43 (Ma et al., 2007; RETRACTION, 2016). Although such a role seems to be supported by the observed ∼6-fold increased accumulation of the RC47 intermediate in the *lpa2* mutant compared to the wild type (Figure 4; Table 2), there are more arguments against it: first, a block of PSII assembly at the level of RC47 would be expected to result in the accumulation of early assembly intermediates, i.e., RC (D1-PsbI:D2-PsbE/F), D1-PsbI, D2-PsbE/F, free CP43 and CP47 – neither of them was observed. The lack of the early assembly intermediates cannot be due to a problem with the synthesis of D1, D2, and CP47, as their synthesis rates and the stability of the newly made proteins were unaffected in the *lpa2* mutant (Figure 2B). Also, the D2 acceptor Cyt *b*_*559*_ (PsbE/F) accumulated to very high levels in the *lpa2* mutant (the D1 acceptor PsbI escaped detection by mass spectrometry). Second, the accumulation of PSII monomers was comparably mildly affected in the *lpa2* mutant (monomers accumulated to 60% of wild type levels), while dimers (26% of wild type levels) and supercomplexes (7% of wild type levels) were much more affected (Figure 4; Table 2). If PSII assembly was impaired at the level of CP43 assembly into RC47, we would have expected monomer assembly to be much more affected.

We therefore propose that LPA2 acts at a step during PSII assembly without which PSII monomers and especially further assemblies become intrinsically unstable and prone to degradation. More prone to degradation are the chlorophyll-binding proteins CP43, CP47, D1, and D2 which therefore are the least abundant PSII core subunits in the *lpa2* mutant. Particularly sensitive is CP43, as it accumulated to lowest levels of all PSII subunits in the mutant (Table 1). PsbE/F, PsbJ, PsbL, and subunits of the oxygen evolving complex are minor targets of degradation and therefore can accumulate as unassembled proteins. The following observations support this idea: first, a normal assembly into monomers would explain why synthesis rates and stability of newly made D1, D2, CP43, and CP47 are not impaired in the *lpa2* mutant. Second, in pulse-chase experiments combined with BN-PAGE, Ma et al. (2007); (RETRACTION, 2016) observed a normal chase of radioactivity into RC47 in the *lpa2* mutant after 15 min but reduced chase into PSII monomers, dimers and supercomplexes compared to the wild type. However, after 60 min of chase, more radioactivity went into PSII monomers and RC47 in the mutant than in the wild type, while little went into PSII dimers and supercomplexes. It appears as if newly made PSII monomers would bounce back after the attempt to assemble into dimers and supercomplexes. Third, the comigration of pheophorbide a oxygenase PAO5 with RC47 only in the mutant is indicative for an active degradation of chlorophyll within RC47 in the thylakoid membrane. Hence, the RC47 accumulating in the *lpa2* mutant would largely be a degradation product rather than an assembly intermediate. At least in *Synechocystis*, both have been proposed to share the same composition (Boehm et al., 2012). Not in support of the idea of increased degradation of PSII cores is that we found no obvious change in the abundance or migration profiles of potentially involved major proteases in or at the thylakoid membranes, including FTSH1/2, DEG1C, and DEG1B (Figure 3B) (Malnoe et al., 2014; Theis et al., 2019). However, the capacity of the constitutively expressed proteases might be sufficient for this task.

At which step of PSII assembly could LPA2 be involved? One possibility is that LPA2 is involved in the efficient assembly of a small PSII core subunit, for example PsbI. In the *Chlamydomonas mbi1* mutant lacking PsbI, the acceptor of newly made D1, PSII assembly proceeded up to the monomer stage, but not beyond (Wang et al., 2015). And in *Synechocystis* the absence of PsbI resulted in a destabilization of CP43 binding within PSII monomers and dimers (Dobakova et al., 2007). PsbJ also appeared an attractive candidate, as it was the only PSII core subunit identified that accumulated to higher levels in the *lpa2* mutant than in the wild type (Table 1; Figure 4). However, we could detect PsbJ in monomers, dimers and supercomplexes in the mutant (Table 2). Unfortunately, mass spectrometry was blind for all other small PSII core subunits. Another possibility is that the *lpa2* mutant is impaired in the C-terminal processing of D1 because the absence of LPA2 came along with the absence of LPA19 (Figure 3B), which facilitates this step in *Arabidopsis* (Wei et al., 2010). LPA19 was detected with three peptides with average identification scores and low ion intensities in all three wild-type replicates (Supplemental Dataset 3), therefore it appears unlikely that we simply missed the protein in the mutant. The *Arabidopsis lpa19* mutant resembles the phenotype of the *Chlamydomonas lpa2* mutant regarding the lower accumulation of PSII core subunits, reduced PSII maximum quantum efficiency, and sensitivity to high light intensities (Wei et al., 2010). Clearly, more work is required to pin down at which step of PSII assembly LPA2 is acting precisely.

### Novel proteins that potentially interact with PSII

Complexome profiling allowed us to identify three novel proteins potentially interacting with PSII. One of them, PBA1, was found to co-migrate with PSII core subunits. Here, the distinct migration profiles exhibited by the core subunits in wild type and *lpa2* mutant increased the confidence of true co-migration and the detection of two peptides the confidence of its identification (Figures 3A and 4; Supplemental Dataset 3). The protein is conserved only in green algae, diatoms, Eustigmatophytes, and brown algae (Figure 5A), which might explain why it has not been detected so far. Its small size (6.4 kDa) and the predicted transmembrane helix are typical features of small PSII core subunits (Pagliano et al., 2013). CGLD16 also is a small protein (7.9 kDa) containing a predicted transmembrane helix. It co-migrated only with PSII monomers and RC47 and, as expected, more with monomers in the wild type, and more with RC47 in the *lpa2* mutant (Figures 3A and 4). Although its detection was based on a single peptide, this was with a high identification score (Supplemental Dataset 3). This protein is conserved in the green lineage and diatoms (Figure 5B) and could potentially represent a novel factor involved in PSII assembly or the regulation of PSII complex dynamics.

The small size of PBA1 and CGLD16 justified that their potential interaction with the different PSII assembly states did not change their migration properties. This is different for the third protein found to potentially interact only with RC47 and only in the *lpa2* mutant, PAO5, which has a predicted molecular mass of 52 kDa and was identified with three peptides of low ion intensity (Supplemental Dataset 3). Its co-migration with RC47 implies that PAO5 has replaced another protein of similar molecular mass in RC47, such as CP47, D1, or D2. Therefore, it could also be that PAO5 interacts with another protein complex and just by chance comigrates with RC47. PAO5 encodes a protein with Rieske-type [2Fe-2S] domain and a pheophorbide a oxygenase domain. It is in the same gene family as the *Arabidopsis* accelerated cell death 1 (ACD1) protein that is involved in chlorophyll a breakdown to avoid chlorophyll phototoxicity (Pružinská et al., 2003; Kuai et al., 2018). However, the closest homologs of PAO5 are cyanobacterial proteins while a close homolog is missing in *Arabidopsis* (Figure 5 and Supplemental Figure 5). Given the 6-fold higher abundance of RC47 in the mutant compared to wild type and the necessity to cope with chlorophyll phototoxicity upon degradation of non-productive PSII assembly intermediates, we believe that the identification of PAO5 only in the *lpa2* mutant is interesting. Clearly, its direct interaction partners need to be identified by co-immunoprecipitation or related approaches.

In this study, we analyzed our complexome profiling dataset with a focus on the major photosynthetic complexes in the thylakoid membranes. However, the dataset is so rich that proteins involved in many other aspects can be investigated, such as complexes of the mitochondrial respiration chain or complexes formed by TCA cycle enzymes (Rugen et al., 2020). These are in the dataset, because tubular mitochondria are closely connected with chloroplasts in *Chlamydomonas* (Niemeyer et al., 2020) and contaminate thylakoid preparations.

## Methods

### Strains and cultivation conditions

*Chlamydomonas reinhardtii* wild-type CC-4533 and *lpa2* mutant strain LMJ.RY0402.141537 from the *Chlamydomonas* library project (Li et al., 2016) were obtained from the *Chlamydomonas* Resource Center. The *lpa2* mutant was used as recipient strain for transformation with plasmid pMBS683 to generate complemented lines c10 and c11, and pMBS684 to generate the complemented line cHA13. Transformation was done via electroporation. Unless indicated otherwise, cultures were grown mixotrophically in TAP medium (Kropat et al., 2011) on a rotatory shaker at 25°C and ∼30 µmol photons m^-2^ s^-1^. Cell densities were determined using a Z2 Coulter Counter (Beckman Coulter).

### Cloning of the construct for complementing the *lpa2* mutant

The *LPA2* coding sequence was amplified by PCR from *Chlamydomonas* cDNA with primers 5′-aagaagAcAGAATGCAGACCTGCTTTTCA-3’ and 5’-ttgaagacttcgaaccCTGCTTCTGGATCTGTCCGGGC-3′ (lower case letters indicate altered bases to introduce BpiI recognition sites). The resulting 552 bp PCR product was cloned into the recipient plasmid pAGM1287 (Weber et al., 2011) by restriction with BbsI and ligation with T4 ligase, resulting in the level 0 construct pMBS680. This level 0 part was then complemented with level 0 parts (pCM) from the *Chlamydomonas* MoClo toolkit (Crozet et al., 2018) to fill the respective positions in level 1 modules as follows: A1-B2 – pCM0-020 (*HSP70A-RBCS2* promoter + 5’UTR); B5 – pCM0-100 (3xHA) or pCM0-101 (MultiStop); B6 – pCM0-119 (*RPL23* 3’UTR). The *HSP70A-RBCS2* fusion promoter used here contains −467 bp of *HSP70A* sequences upstream from the start codon in optimal spacing with respect to the *RBCS2* promoter (Lodha et al., 2008; Strenkert et al., 2013). The level 0 parts and destination vector pICH47742 (Weber et al., 2011) were combined with BsaI and T4 DNA ligase and directionally assembled into level 1 modules pMBS682(3xHA) and pMBS681 (MultiStop), which were then combined with pCM1-01 (level 1 module with the *aadA* gene conferring resistance to spectinomycin) from the *Chlamydomonas* MoClo kit, with plasmid pICH41744 containing the proper end-linker, and with destination vector pAGM4673 (Weber et al., 2011), digested with BbsI, and ligated to yield level 2 devices pMBS684 and pMBS683, respectively.

### RNA extraction and qRT-PCR

10^8^ cells were harvested by centrifugation at 4500 rpm for 2 min, resuspended in lysis buffer (0.6 M NaCl, 0.1 M Tris-HCl, pH 8, 10 mM EDTA, 4 % SDS), snap-frozen in liquid N_2_ and stored at − 80 °C until further use. For RNA extraction, samples were incubated at 65 °C for 10 min, supplemented with 2 M KCl, incubated for 15 min on ice and centrifuged at 12,500 rpm for 15 min. Two extraction steps followed with phenol/chloroform/isoamylalcohol (25:24:1) and chloroform/isoamylalcohol (24:1) and RNA was precipitated over night at 4 °C in 8 M LiCl. The precipitate was collected by centrifugation at 12,500 rpm for 15 min, resuspended in RNase free H_2_O and supplemented with 3 M Na-acetate, pH 5.2. The RNA was precipitated for 45 min in 100 % ethanol on ice and washed with 70 % ethanol. After a further centrifugation step, the pelleted and dried RNA was resuspended in RNase free H_2_O, the concentration was determined spectrophotometrically using a NanoDrop 2000 (Thermo Scientific) and the RNA quality was confirmed using agarose gel electrophoresis. DNA contaminations were removed via RNase-free Turbo DNase (Ambion) and complementary DNA (cDNA) synthesis was performed using the M-MLV reverse transcriptase (Promega), deoxynucleotide triphosphates, and oligo-d(T)_18_ primers. Quantitative reverse transcription-PCR (qRT-PCR) was performed using the StepOnePlus RT-PCR system (Applied Biosystems) and the 5x HOT FIREPol® EvaGreen® qPCR Supermix kit from Solis BioDyne. Each reaction contained the manufacturer’s master mix, 150 nM of each primer and cDNA corresponding to 10 ng of input RNA in the reverse transcription reaction. The reaction conditions were as follows: 95 °C for 10 min, followed by cycles of 95 °C for 15 s, 65 °C for 20 s and 72 °C for 20 s, up to a total of 40 cycles. Primers used for the amplification of the *LPA2* transcript were 5’-GGGCTTTGGTTCAGAGACGG-3’ and 5’-TGCGTTCACCTTGACCTTGG-3’. Primers used for the amplification of the endogenous control (*CBLP2*) were 5’-GCCACACCGAGTGGGTGTCGTGCG-3’ and 5’-CCTTGCCGCCCGAGGCGCACAGCG-3’.

### Protein analyses

Cells were harvested by centrifugation, resuspended in sample buffer containing 75 mM Tris-HCl, pH 6.8, 2 % (w/v) SDS and 10 % (v/v) glycerol, boiled at 95 °C and centrifuged. After quantification of protein concentrations according to (Bradford, 1976), Laemmli buffer (Laemmli, 1970) was added to samples and SDS-PAGE and semi-dry western blotting were performed as described previously (Liu et al., 2005). Antisera used were against D1 (Agrisera AS05 084), CP43 (Agrisera AS11 1787), CP47 (Agrisera AS04 038), PsaA (Agrisera AS06 172), PSAN (M. Schroda, unpublished data), CytF (Pierre and Popot, 1993), CGE1 (Schroda et al., 2001), LHCBM9 (M. Schroda, unpublished data), CF1β (Lemaire and Wollman, 1989), RPL1 (Ries et al., 2017) and the HA-tag (Sigma-Aldrich H3663). Anti-rabbit-HRP (Sigma-Aldrich) and anti-mouse-HRP (Santa Cruz Biotechnology sc-2031) have been used as secondary antibodies.

BN-PAGE with whole cell proteins was carried out as described previously (Schagger and von Jagow, 1991) with minor modifications. Briefly, 10^8^ cells were pelleted, washed with 750 µl of TMK buffer (10 mM Tris-HCl, pH 6.8, 10 mM MgCl_2_, 20 mM KCl) and resuspended in 350 µl of ACA buffer (750 mM ε-aminocaproic acid, 50 mM Bis-Tris/HCl, pH 7.0, 0.5 mM EDTA) supplemented with 0.25x protease inhibitor (Roche). Cells were then broken by sonication and starch and cell debris were removed by centrifugation at 300 g for 5 min at 4 °C. Cell lysates (equivalent to 0.8 µg/µl of protein) were solubilized with 1 % α-DDM for 20 min on ice and in darkness. Insolubilized material was pelleted at 13,200 rpm for 10 min at 4 °C and the supernatant was supplemented with loading buffer (0.5 M ε-aminocaproic acid, 75 % glycerol, 2.5 % Serva Blue G-250 (Roth)). After three cycles of centrifugation at 13,200 rpm for 10 min at 4 °C and transferring the supernatant to a fresh tube, samples were loaded on a 4-15 % BN acrylamide gel.

The isolation of thylakoids was performed according to (Chua and Bennoun, 1975) with minor modifications. Briefly, ∼2 * 10^9^ cells were pelleted and washed with 25 mM HEPES-KOH, pH 7.5, 5 mM MgCl_2_ and 0.3 M sucrose, before resuspending in the same buffer supplemented with protease inhibitor (Roche). Cells were then lysed using a BioNebulizer (Glas-Col) with an operating N_2_ pressure of 1.5-2 bar. After centrifugation at 5000 rpm for 10 min, the pellet was washed with 5 mM HEPES-KOH, pH 7.5, 1 mM EDTA and 0.3 M sucrose before resuspending in 5 mM HEPES-KOH, pH 7.5, 1 mM EDTA and 1.8 M sucrose. After a sucrose step gradient (0.5 M, 1.3 M, 1.8 M) centrifugation at 24,000 rpm for 1 h, intact thylakoids, floating between the 1.3 M and 1.8 M layers, were collected and diluted with 5 mM HEPES-KOH, pH 7.5 and 1 mM EDTA. After another centrifugation at 13,000 rpm for 1 h, the pellet was resuspended in small volumes of the same buffer.

For pulse labelling experiments, cells in the exponential growth phase (2 ×10^6^ cells ml^-1^) from a 100-ml culture were harvested by centrifugation, washed with minimum medium and re-suspended in 1/20th volume of minimum medium. Cells were allowed to recover and to deplete their intracellular carbon pool for 1.5 hours under dim light (20 µE m^-2^ s^-1^) and strong aeration at 25°C. 10 µM cycloheximide and 10 µCi ml^-1^ Na-^14^C acetate (PerkinElmer: 56.6 mCi mM^-1^) were then added to the culture. After 7 min the pulse was stopped by transferring the cells into 35 ml of ice-cold TAP medium containing 50 mM non-radioactive acetate. Cell samples were collected immediately and after incubation for 20 and 60 min (chase) by centrifugation, resuspended in 0.1 M DTT and 0.1 M Na_2_CO_3_, frozen in liquid nitrogen, and kept at – 80°C until analysis.

### In-gel digestion and mass spectrometry

Coomassie stained BN-PAGE gel pieces were destained by repeated cycles of washing with 40 mM NH_4_HCO_3_ for 5 min and incubating in 70 % acetonitrile for 15 min, until they were colorless. They were then dehydrated completely by adding 100 % acetonitrile for 5 min and dried under vacuum. Samples were then digested by covering the gel pieces in 10 ng/µl trypsin in 40 mM NH_4_HCO_3_ and incubating them over night at 37 °C, before first, hydrophilic peptides were extracted with 10 % acetonitrile and 2 % formic acid for 20 min and afterwards, all other tryptic peptides were extracted with 60 % acetonitrile and 1 % formic acid. Samples were then desalted according to (Rappsilber et al., 2007). Mass spectrometry was performed basically as described previously (Hammel et al., 2018). For peptide separation, a HPLC flow rate of 4 μl/min and 21 min gradients from 2 % to 33 % HPLC buffer B were employed (buffer A 2% acetonitrile, 0.1% formic acid; buffer B 90% acetonitrile, 0.1% formic acid). MS1 spectra (350 m/z to 1250 m/z) were recorded for 250 ms and 25 MS/MS scans (100 m/z to 1500 m/z) were triggered in high sensitivity mode with a dwell time of 50 ms resulting in a total cycle time of 1550 ms. Analyzed precursors were excluded for 5 s, singly charged precursors or precursors with a response below 500 cps were excluded completely from MS/MS analysis.

### Evaluation of MS data

The analysis of MS runs was performed using MaxQuant version 1.6.0.1 (Cox and Mann, 2008). Library generation for peptide spectrum matching was based on *Chlamydomonas reinhardtii* genome release 5.5 (Merchant et al., 2007) including chloroplast and mitochondrial proteins. Oxidation of methionine and acetylation of the N-terminus were considered as peptide modifications. Maximal missed cleavages were set to 3 and peptide length to 6 amino acids, the maximal mass to 6000 Da. Thresholds for peptide spectrum matching and protein identification were set by a false discovery rate (FDR) of 0.01. The mass spectrometry data have been deposited to the ProteomeXchange Consortium via the PRIDE partner repository with the dataset identifier XXX (*in work*). Total protein group intensities varied between samples. For sample normalization, the total ion intensity sum (TIS) of every protein and gel slice was calculated for each of the six samples (3 x wild type and 3 x mutant). For every sample, a correction factor (CF) was determined by dividing every TIS by the average total intensity sum. Subsequently, every intensity value was divided by its corresponding correction factor, to equalize all TISs. For further analysis, proteins identified by non-proteotypic peptides were discarded. Protein identifiers were annotated with MapMan ontology terms, Gene Ontology (GO) terms, and proposed subcellular localization using the Functional Annotator (FATool available at http://iomiqsweb1.bio.uni-kl.de:8015/). A Welch test was performed for each protein by considering the sums of all 36 normalized slice intensities for each sample and testing three wild type sums against three mutant sums. The distance of the average migration profiles for every protein was calculated as the Euclidean distance between wild type and mutant. To adjust for amplitude-introduced bias, each distance was divided by the maximal average intensity of wild type or mutant, respectively. Data normalization and analysis were performed using FSharp.Stats (https://github.com/CSBiology/FSharp.Stats). The migration profiles were visualized using Plotly.NET (https://github.com/muehlhaus/FSharp.Plotly) and the NOVA software (Giese et al., 2014).

### Cell fractionation

Crude fractionation was performed to separate soluble and membrane proteins. 2 * 10^7^ cells were pelleted and resuspended in 1 ml of lysis buffer (10 mM Tris-HCl, pH 8.0, 1 mM EDTA) including 0.25x protease inhibitor (Roche)). A 200 µl whole cell aliquot was taken and supplemented with 50 µl of sample buffer (225 mM Tris-HCl, pH 6.8, 50 % glycerol, 5 % SDS, 0.25 M DTT, 0.05 % bromophenol blue). The remaining solution was frozen in liquid nitrogen and thawed at room temperature for four cycles. After centrifugation at 21,000 g for 20 min, the supernatant, containing soluble proteins, was collected. The pellet fraction was resuspended in lysis buffer. Sample buffer was added to both protein extracts prior to boiling at 95 °C and protein separation via SDS-PAGE.

### Chlorophyll fluorescence measurements

Maximum quantum efficiency of PSII (F_v_/F_m_) was measured using a pulse amplitude-modulated Mini-PAM fluorometer (Mini-PAM, H. Walz, Effeltrich, Germany) essentially according to the manufacturer’s protocol after 3 min of dark adaptation (1 s saturating pulse of 6000 μmol photons m^-2^ s^-1^, gain = 4).

### Sequence alignments and predictions

Putative chloroplast transit peptides of LPA2 homologs were predicted with ChloroP (Emanuelsson et al., 1999) and putative transmembrane helices with HMMTOP (Tusnady and Simon, 2001). Sequence alignments were done with CLUSTALW (https://www.genome.jp/tools-bin/clustalw) and displayed with GeneDoc.

## Acknowledgements

We would like to thank Peter Kroth for helping with the annotation of bipartite transit peptides from heterokonts.

